# Heterologous Expression of Human Norovirus GII.4 VP1 Leads to Assembly of T=4 Virus-Like Particles

**DOI:** 10.1101/583385

**Authors:** Jessica Devant, Götz Hofhaus, David Bhella, Grant S. Hansman

## Abstract

Human noroviruses are a leading cause of acute gastroenteritis, yet there are still no vaccines or antivirals available. Expression of the norovirus capsid protein (VP1) in insect cells typically results in the formation of virus-like particles (VLPs) that are morphologically and antigenically comparable to native virions. Previous structural analysis of norovirus VLPs showed that the capsid has a T=3 icosahedral symmetry and is composed of 180 copies of VP1 that are folded into three quasi-equivalent subunits (A, B, and C). In this study, we determined the cryo-EM VLP structures of two GII.4 variants, termed CHDC-1974 and NSW-2012. Surprisingly, we found that greater than 95% of these GII.4 VLPs were larger than virions and 3D reconstruction showed that these VLPs exhibited T=4 icosahedral symmetry. We found that the T=4 VLPs showed several structural differences to the T=3 VLPs. The T=4 particles assemble from 240 copies of VP1 that adopt four quasi-equivalent conformations (A, B, C, and D) that form two distinct dimers, A/B and C/D. The T=4 protruding domains were elevated ∼21-Å off the capsid shell, which was ∼7-Å more than the previously studied GII.10 T=3 VLPs. A small cavity and flap-like structure at the icosahedral twofold axis disrupted the contiguous T=4 shell, a consequence of the D-subunit S-domains having smaller contact interfaces with neighboring dimers. Overall, our findings that old and new GII.4 VP1 sequences assemble T=4 VLPs might have implications for the design of potential future vaccines.

**IMPORTANCE:** The discovery that the GII.4 VLPs have a T=4 symmetry is of significance, since this represents the first known T=4 calicivirus structure. Interestingly, the GII.4 2012 variant shares 96% amino acid identity with a current GII.4 VLP vaccine candidate sequence, which suggests that this vaccine might also have a T=4 symmetry. Our previous results with these GII.4 VLPs showed functional binding properties to antibodies and Nanobodies that were raised against T=3 (GII.10) VLPs. This suggests that the T=4 VLPs were antigenically comparable to T=3 particles, despite the obvious structural and size differences. On the other hand, these larger T=4 VLPs with novel structural features and possibly new epitopes might elicit antibodies that do not recognize equivalent epitopes on the T=3 VLPs. Further structural and binding studies using a library of GII.4-specific Nanobodies are planned in order to precisely investigate whether new epitopes are formed.

## INTRODUCTION

Human noroviruses are members of the *Caliciviridae* family and are a leading cause of outbreaks of acute gastroenteritis. The virus has a positive sense, single stranded RNA genome of ∼7.7 kbp. The genome is organized into three open reading frames (ORFs), where ORF1 encodes nonstructural proteins and ORF2 and ORF3 encode a major structural protein (termed VP1) and a minor structural protein (termed VP2), respectively. Noroviruses are genetically diverse and based on VP1 sequences have been classified into seven genogroups (GI-GVII), where GI, GII, and GIV cause infections in humans (1, 2). The GI and GII are further subdivided into numerous genotypes, with GII genotype 4 (GII.4) recognized as the most prevalent and clinically important strain (3, 4).

Recently, two human norovirus cell culture systems were developed (5, 6). However, mechanistic studies of norovirus structure and biology, such as interaction with the host receptor(s), remain challenging owing to difficulties in preparing large-scale virion preparations. Expression of ORF2 alone in insect or mammalian cells can result in the formation of virus-like particles (VLPs) that are antigenically and morphologically similar to native virions (2, 7-10). Expression of norovirus VLPs has permitted studies on host binding factors, interactions with norovirus-specific antibodies, and structural studies (2, 11-16). Indeed, histo-blood group antigens (HBGAs) and bile acids were shown to be important binding co-factors for human norovirus VLPs and/ or virions (17-22).

Structural analysis of GI.1 VLPs revealed that VP1 is separated into two distinct domains: a shell domain (S domain) that encloses the RNA and a protruding domain (P domain) that binds to co-factors, such as HGBAs and bile acids (8, 19, 21). A hinge region, which is typically composed of 10-14 amino acids, connects the S and P domains. The P domain has a β-barrel fold that is structurally conserved in the *Caliciviridae* family. Dimerization of the P domains forms arch shaped protrusions that are visible in electron microscopy images. The P domain is further subdivided into P1 and P2 subdomains, where the P2 subdomain is an insertion in the P1 subdomain and is the most variable region on the capsid (8).

Structural studies have shown that caliciviruses have a common overall organization of T=3 icosahedral symmetry and are comprised of 180 copies of VP1 (7, 8, 23-25). Within the asymmetric unit, VP1 adopts three quasi-equivalent conformations, termed A, B, and C (8). The norovirus A and B subunits assemble into 60 dimers (termed A/B) at the quasi twofold axis, whereas the C subunits assemble into 30 C/C dimers that are located at the strict icosahedral twofold axis. For the GI.1 VLPs, the A/B dimers have a convex S domain conformation, whereas the C/C dimers have a flat S domain conformation (8). The conformational differences within these dimers likely facilitates the curvature of the virus particle to form a closed shell, such features are commonly seen in other T=3 icosahedral viruses (26). Interestingly, smaller norovirus VLPs (∼25 nm in diameter) that are assumed to have a T=1 icosahedral symmetry have also been reported (22, 27); however to date, structures of these smaller VLPs have not been published.

Here, we show the cryo-EM structures of VLPs for two GII.4 variants that were identified in 1974 and 2012, termed CHDC-1974 and NSW-2012, respectively (15, 28). We show that these VLPs have a T=4 icosahedral symmetry and are composed of 240 copies of VP1. In order to form the T=4 icosahedral structure, VP1 adopts four quasi-equivalent conformations, termed A, B, C, and D, giving rise to two distinct types of dimers, termed A/B and C/D. The VLPs consisted of 60 A/B dimers and 60 C/D dimers, with B, C, and D subunits located at the twofold axis, and the A subunit at the fivefold axis. As GII.4 VLPs are currently under consideration as vaccine candidates, our findings might have implications with respect to the likely antigenic properties of such assemblies.

## MATERIALS AND METHODS

### VLP and virion preparation

The NSW-2012 and CHDC-1974 VLPs (Genebank accession numbers JX459908 and ACT76142, respectively) were expressed in a baculovirus system as previously described (29-31). Briefly, the bacmid containing the recombinant VP1 gene was transfected in Sf9 insect cells. After incubation for five days, the culture medium was centrifuged for 10 min at 3,000 rpm at 4°C. The recovered baculovirus was subsequently used to infect Hi5 insect cells. At five days post infection, the culture medium was centrifuged for 10 min at 3,000 rpm at 4°C and then 1 h at 6,500 rpm at 4°C. The VLPs in the supernatant were concentrated by ultracentrifugation at 35,000 rpm (Beckman Ti45) for 2 h at 4°C and then further purified using CsCl equilibrium gradient ultracentrifugation at 35,000 rpm (Beckman SW56) for 18 h at 4°C. To remove the CsCl, the VLPs were pelleted for 2 h at 40,000 rpm (Beckman TLA55) at 4°C and subsequently resuspended in PBS (pH 7.4). GII.4 virions in stool were also purified using this centrifugation technique, except for the CsCl gradient step.

### Negative stain electron microscopy

The integrity of the VLPs was confirmed using negative stain electron microscopy (EM). The VLPs were diluted 1:30 in distilled water and applied to EM grids. The grids were washed with distilled water, stained with 0.75% uranyl acetate, and the excess uranyl acetate was removed with filter paper. Virion samples were applied to EM grids, washed with water, fixed with 4% glutaraldehyde, and then stained as above. EM images were acquired on a Zeiss 910 electron microscope at 50,000× magnification.

### Cryo-EM data sample preparation and data collection

UNSW-2012 and CHDC-1974 VLPs (3 µl) were applied to freshly glow discharged Quantifoil holey carbon support films (R1.2/1.3) and blotted for 18 seconds at 100% humidity and 10°C before being plunged in liquid ethane using an FEI Mark IV Vitrobot (Thermo Fischer Scientific). Vitrified specimens were imaged on a Titan Krios microscope (Thermo Fisher scientific) operated at 300 keV. NSW-2012 micrographs were acquired with a K2 direct electron detector with Latitude S software (Gatan) at 64,000× magnification corresponding to a pixel size of 2.27 Å/px, while CHDC-1974 micrographs were collected using a K3 direct electron detector at 64,000× magnification, corresponding to a pixel size of 1.375 Å/px.

### Cryo-EM data processing

Initially, the movies containing 16 frames for NSW-2012 and 40 frames for CHDC-1974 were motion corrected using motioncor2 software (32) and defocus estimation was performed using ctffind 4.1 software (33). All further image-processing steps were performed using Relion 2.1 software for NSW-2012 and cryoSPARC software for CHDC-1974 (34, 35). An initial set of 1,000 particles was manually picked for 2D classification to produce averages suitable as references for automated particle picking. The autopicked particles were sorted in a 2D classification step and the best particles were used for calculation of an initial starting model, followed by 3D classification. A subset of particles that generated the highest resolution was selected for further refinement. The 3D refinement and post-processing of NSW-2012 from 10,548 T=4 particles produced a final map at 7.3-Å resolution with icosahedral (I2) symmetry imposed (0.143 FSC cutoff). Smaller NSW-2012 T=3 particles were manually picked and sorted in 2D classification. In total, 391 particles were selected for further 3D structure determination. Refinement of these smaller NSW-2012 T=3 particles yielded a map of 15-Å resolution. For CHDC-1974, a subset of 42,485 particles for refinement led to the calculation of a map at 6.1-Å resolution using the 0.143 FSC cutoff. Cryo-EM T=4 VLP structures for CHDC-1974 (accession number: EMD-4549) and NSW-2012 (EMD-4550) were deposited at EMDB. The cryo-EM VLP structure for NSW-2012 with T=3 icosahedral symmetry is available on request.

### Fitting of the X-ray structures into the density maps

Crystal structures of NSW-2012 P domain (4OOS) and CHDC-1974 P domain (5IYN) were fitted into the respective densities using the “fit in map” command in the UCSF Chimera software (36). Since a high-resolution GII.4 shell domain structure was unavailable, the GI.1 Norwalk virus S domain was extracted from the X-ray crystal structure (1IHM) and fitted into the GII.4 cryo-EM densities using UCSF Chimera software.

## RESULTS AND DISCUSSION

The purpose of this study was to analyze GII.4 VLP architecture and compare these assemblies to the previously solved GI.1 and GII.10 VLP structures. The NSW-2012 VP1 sequence had a single amino acid insertion at position ∼394 (NSW-2012 numbering) compared to CHDC-1974 (Fig. 1). Overall, NSW-2012 and CHDC-1974 shared 89% amino acid identity, with most (45 of 54) amino acid substitutions occurring in the P domain. Negative stain EM images revealed that the VLPs exhibited characteristic norovirus morphology (Fig. 2). However, the diameter of these VLPs was measured to be ∼52 nm, which suggested that the GII.4 VLPs were larger than GII.10 and GI.1 VLPs that had diameters of ∼43 nm and ∼38 nm, respectively.

**Figure 1.**
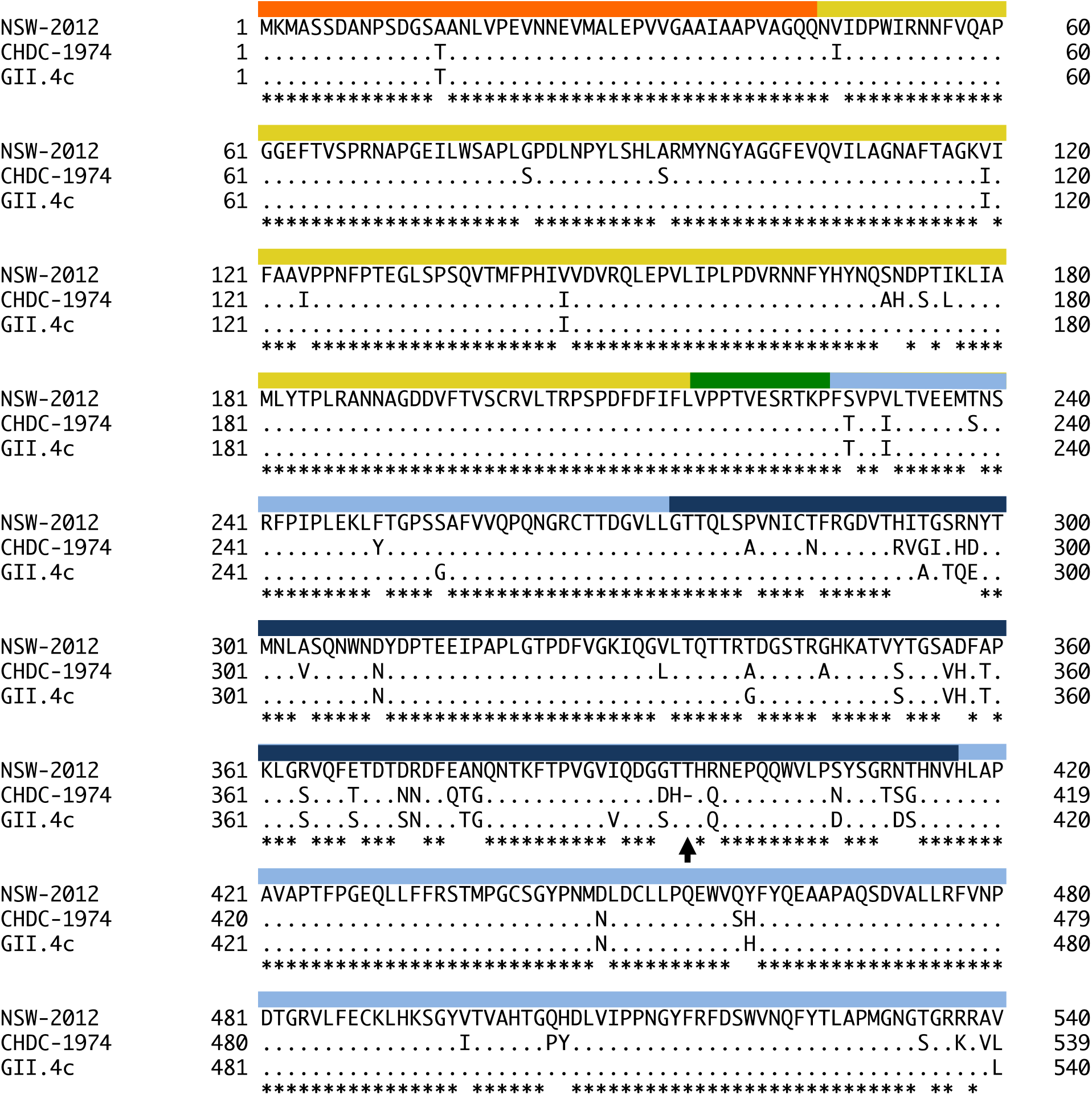
Amino acid sequence alignment of GII.4 VP1. NSW-2012 (JX459908), CHDC-1974 (ACT76142), and GII.4c (42) VP1 amino acid sequences were aligned using ClustalX. The S domain (orange), hinge region (green), P1 subdomain (light blue), and P2 subdomain (navy) were labeled accordingly. Compared to CHDC-1974, NSW-2012 and GII.4c VP1 had a single amino acid insertion (arrow) at position 394 (NSW-2012 numbering). The S domain and hinge region were mainly conserved, whereas most amino acid substitutions were located in the P2 subdomain.

**Figure 2.**
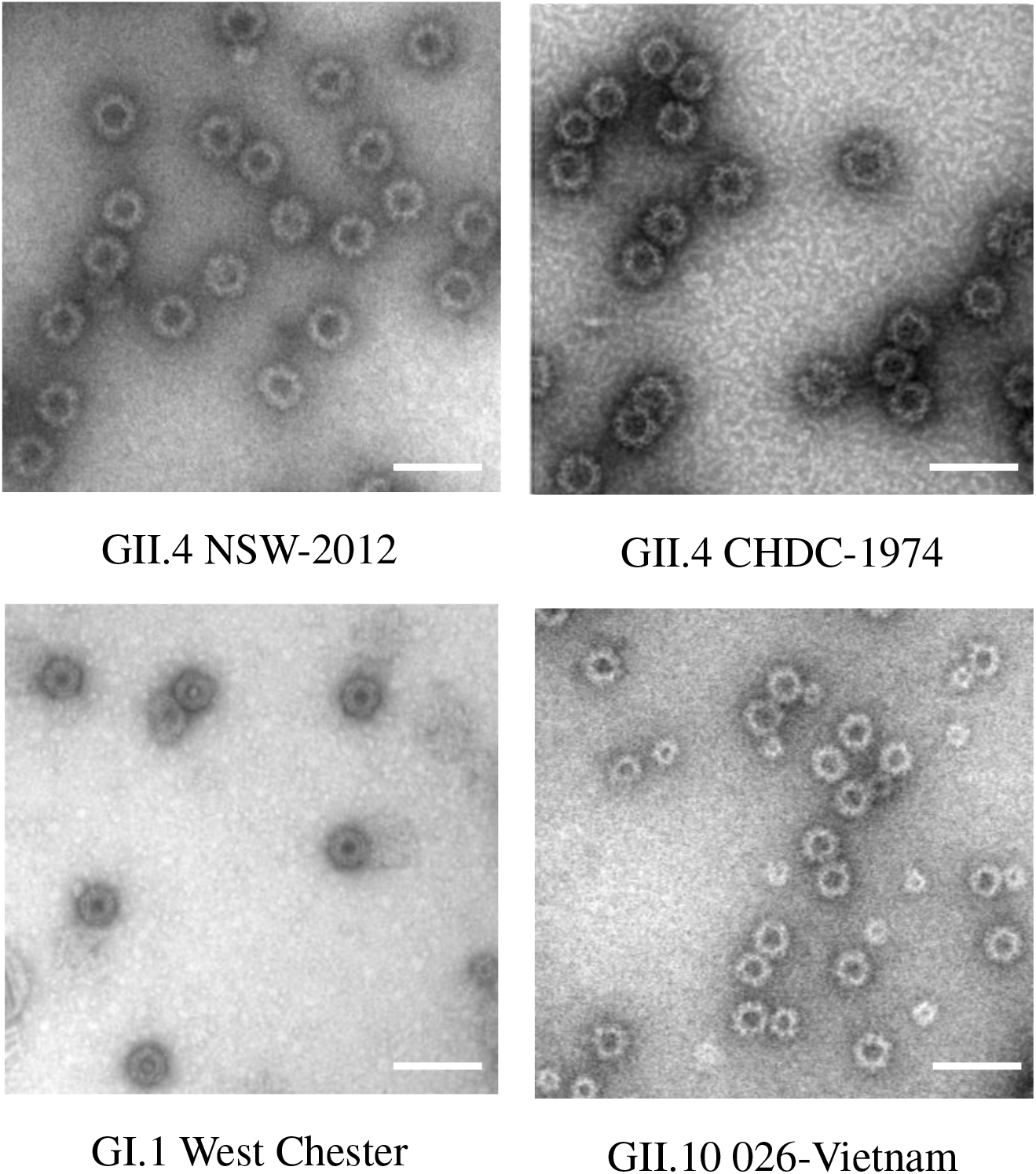
EM images of GI and GII VLPs. Negative stain EM images of norovirus VLPs show the characteristic norovirus virion morphology (50,000× magnification). The diameter of the CHDC-1976 and NSW-2012 VLPs was ∼52 nm, whereas the diameter of GI.1 West Chester (AY502016.1) and GII.10 Vietnam026 (AF504671) VLPs were ∼38 nm and ∼43 nm, respectively (45, 46). The bar represents 100 nm.

### Cryo-EM structure of NSW-2012 T=4 VLPs

The structure of the NSW-2012 VLPs was determined using cryo-EM and 3D icosahedral image reconstruction. The VLPs were mono-disperse in vitreous ice and appeared mostly homogenous in size (Fig. 3A). From the 364 images, 10,548 particles were used for image reconstruction and refined to 7.3-Å resolution (Fig. 3C). Unexpectedly, NSW-2012 VLPs were discovered to have T=4 icosahedral symmetry (Fig. 4). Our data revealed that these VLPs were composed of 240 copies of VP1, rather than 180 VP1 copies as in GI.1 and GII.10 VLPs. The inner diameter of NSW-2012 shell was measured as 32 nm, while the outer capsid diameter was 50 nm. The larger diameter of the T=4 VLPs corresponded to a capsid volume of ∼12,724 nm^3^, which was ∼2.1 times the volume of the GII.10 VLPs that had an inner diameter of 23 nm, corresponding to a volume of ∼5,985 nm^3^ (7).

**Figure 3.**
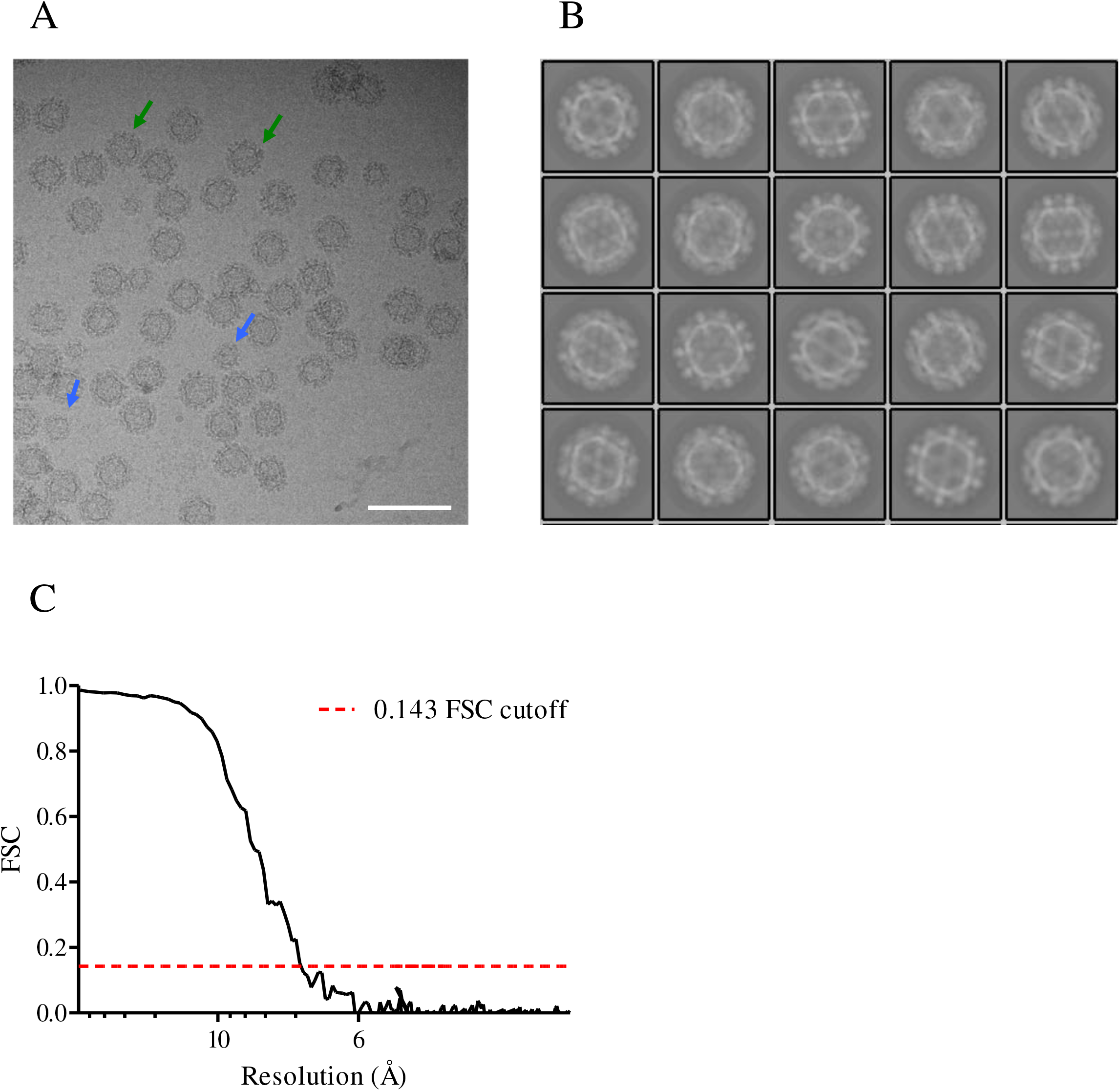
NSW-2012 cryo-EM data processing. (A) A representative cryo-EM micrograph of NSW-2012 VLPs at 64,000× magnification. The blue and green arrows show examples of VLPs measuring ∼46 nm and ∼50 nm, respectively. The scale bar represents 100 nm. B) 2D classification of GII.4 NSW-2012 T=4 VLPs. (C) Gold standard FSC plot of the icosahedral reconstruction of NSW-2012 indicates a resolution of 7.3-Å.

**Figure 4.**
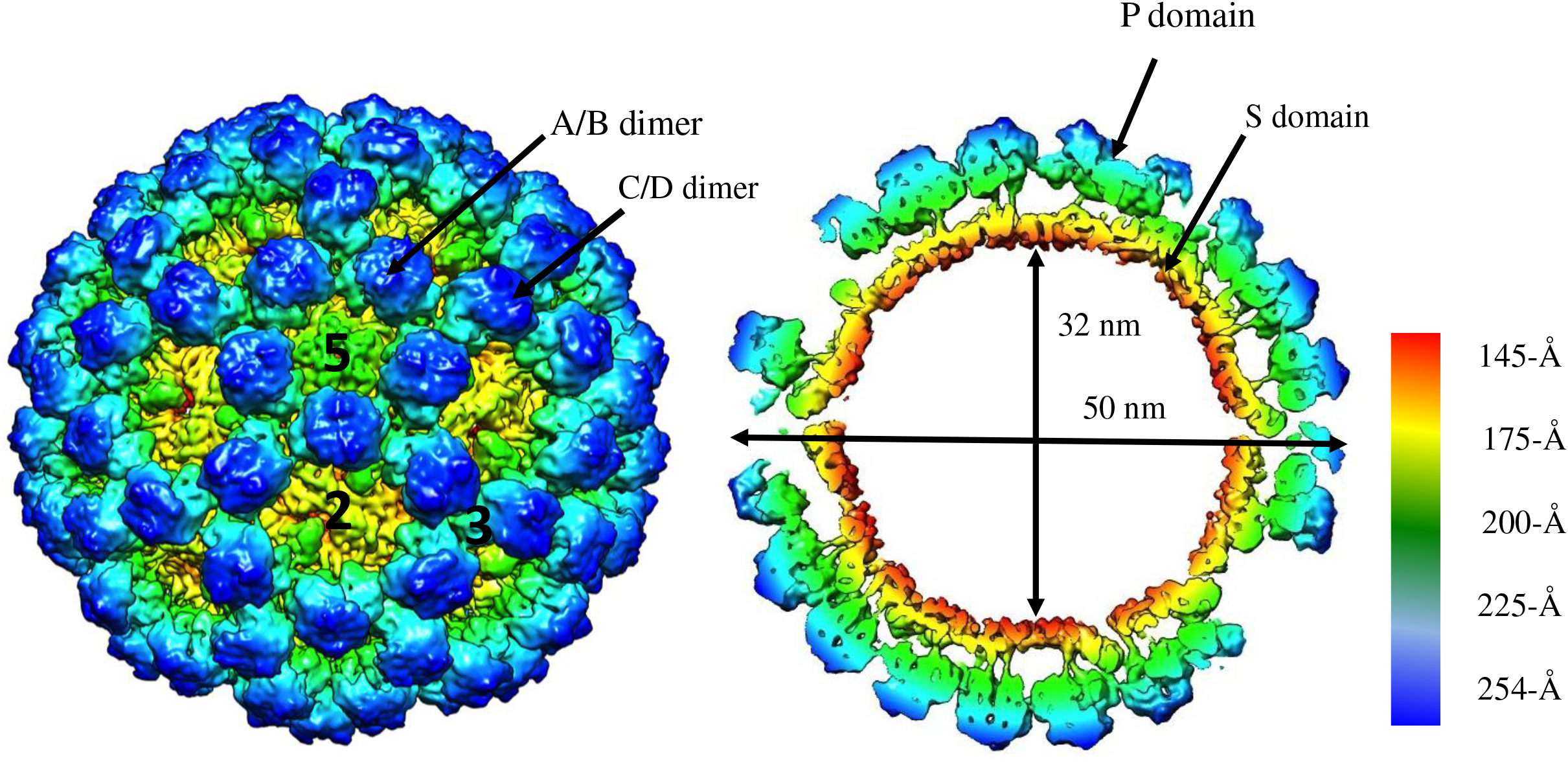
Cryo-EM reconstruction structure of NSW-2012 T=4 VLPs. The left side shows NSW-2012 VLPs have a T=4 icosahedral symmetry (symmetry axis labeled 2, 3, and 5). These VLPs were composed of 240 copies of VP1 and VP1 adapted four quasiequivalent conformations (A, B, C, and D) that gave rise to two distinct dimers (A/B and C/D). At the icosahedral twofold axis, the B, C, and D subunits were alternating, while the A subunits are positioned at the fivefold axis. The right side shows a cutaway section of these VLPs and indicates that the inner and outer diameters are 32 nm and 50 nm, respectively. The P domains are elevated ∼21 Å off the S domain.

Another interesting structural feature of these NSW-2012 VLPs was a small cavity and flap-like structure on the contiguous shell (Fig. 5). This feature was associated with the S domain and found on opposing sides at the twofold axis. This feature may arise as a consequence of the C/D dimer having a stronger curvature, preventing the S-domain from forming the expected contacts with neighboring B and C-type subunits. Alternatively, there might be insufficient space to allow the D subunit S-domain to pack, such that it lies in the plane of the shell without causing steric collision with neighboring subunits.

**Figure 5.**
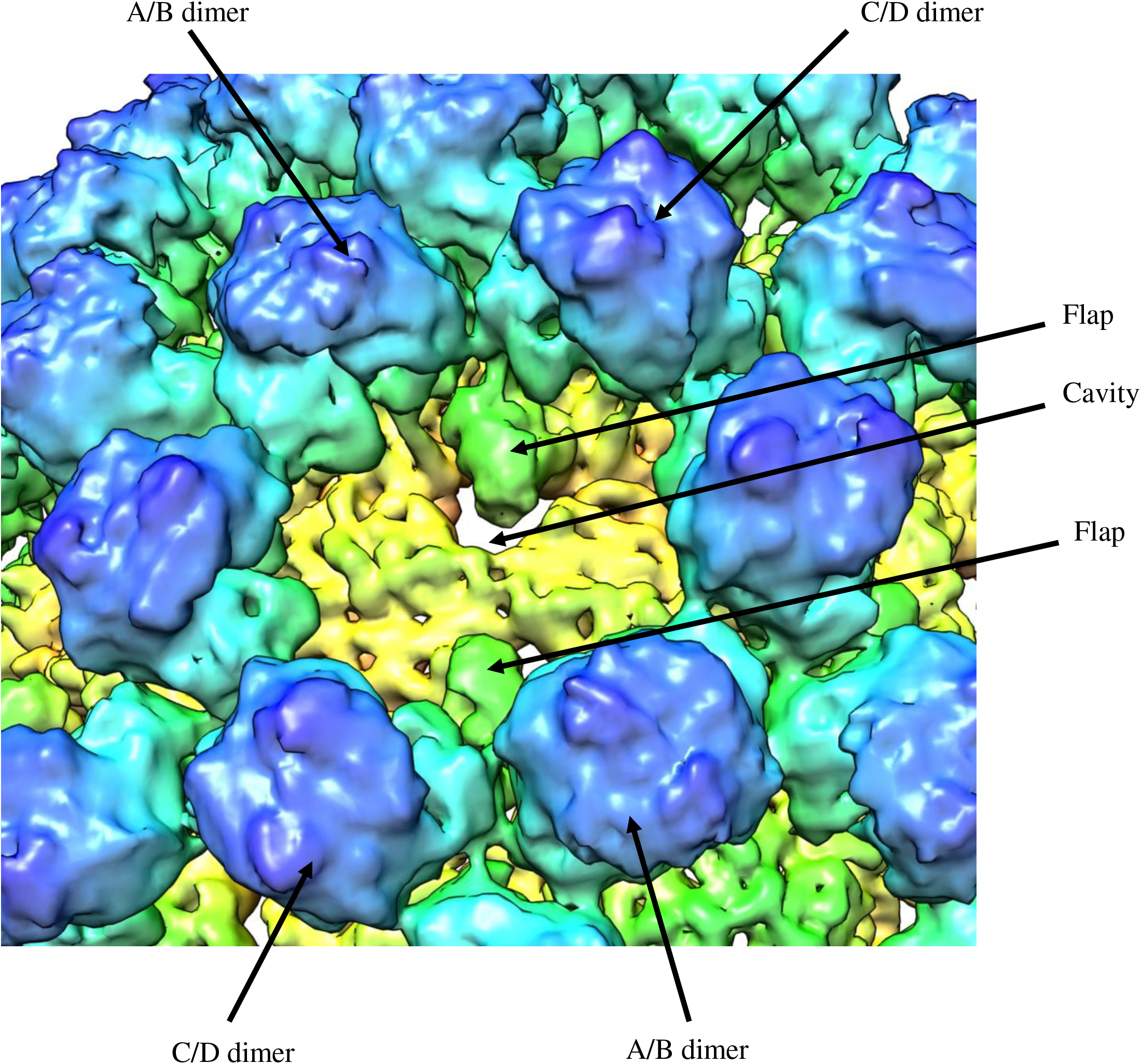
NSW-2012 T=4 VLPs shows several new structural features. The cavity and flap-like structures are observed at the twofold axis and are found on opposing sides. The cavity and flap-like structures are associated with the S domain on the D subunit.

Structural analysis of the T=4 VLPs indicated that VP1 adopted four quasi-equivalent conformations, termed A, B, C, and D. These four subunit conformations formed two distinct dimer classes: A/B and C/D (Figs. 4 and 6). The B, C, and D subunits alternated about the two-fold symmetry axis, while the A subunit was positioned at the fivefold axis (Fig. 4). The T=4 VLPs were composed of 60 A/B dimers and 60 C/D dimers, which was distinct from the T=3 VLPs that have been shown to be assembled from 60 A/B and 30 C/C dimers. We also observed that both A/B and C/D dimers had a convex S domain conformation, which was in contrast to the GI.1 VLPs that consisted of both convex (A/B) and flat (C/C) dimers (8).

**Figure 6.**
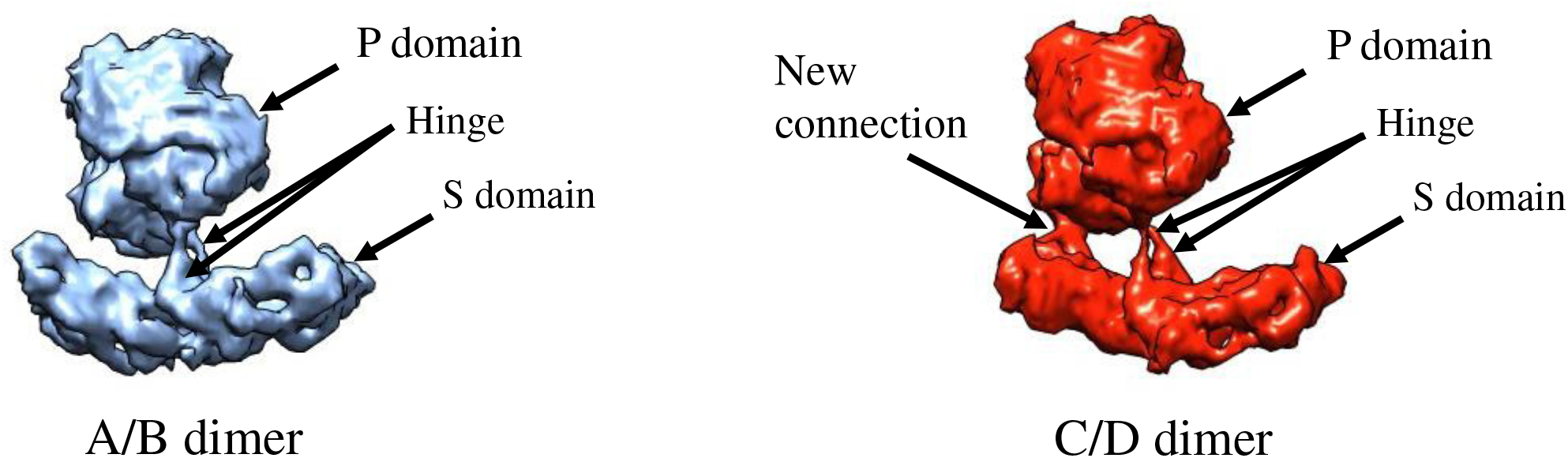
NSW-2012 T=4 VLPs are formed with 60 A/B and 60 C/D VP1 dimers. The A/B and C/D dimers show an equivalent convex conformation of the S domain. An additional connection was also observed between the D subunit of the S and P domain.

In order to better comprehend how VP1 assembled into the T=4 VLPs, the X-ray crystal structure of NSW-2012 P domain (4OOS) and GI.1 S domain (1IHM) were fitted into the VLP density map. We found that the NSW-2012 P domain dimer could be unambiguously positioned into the VLP density, with a cross correlation coefficient of 0.96 (Fig. 7A). This result indicated that the P domain dimers on the T=4 VLPs had not undergone any major structural modifications. In the case of the S domain, the GI.1 S domain needed to be manually positioned into the density map. The GI.1 S domain fitted well into to A/B dimer and the C subunit, while the D subunit needed to be further repositioned (Fig. 7B). This additional fitting in the D subunit was necessary in order to occupy the elevated density of the flap-like regions.

**Figure 7.**
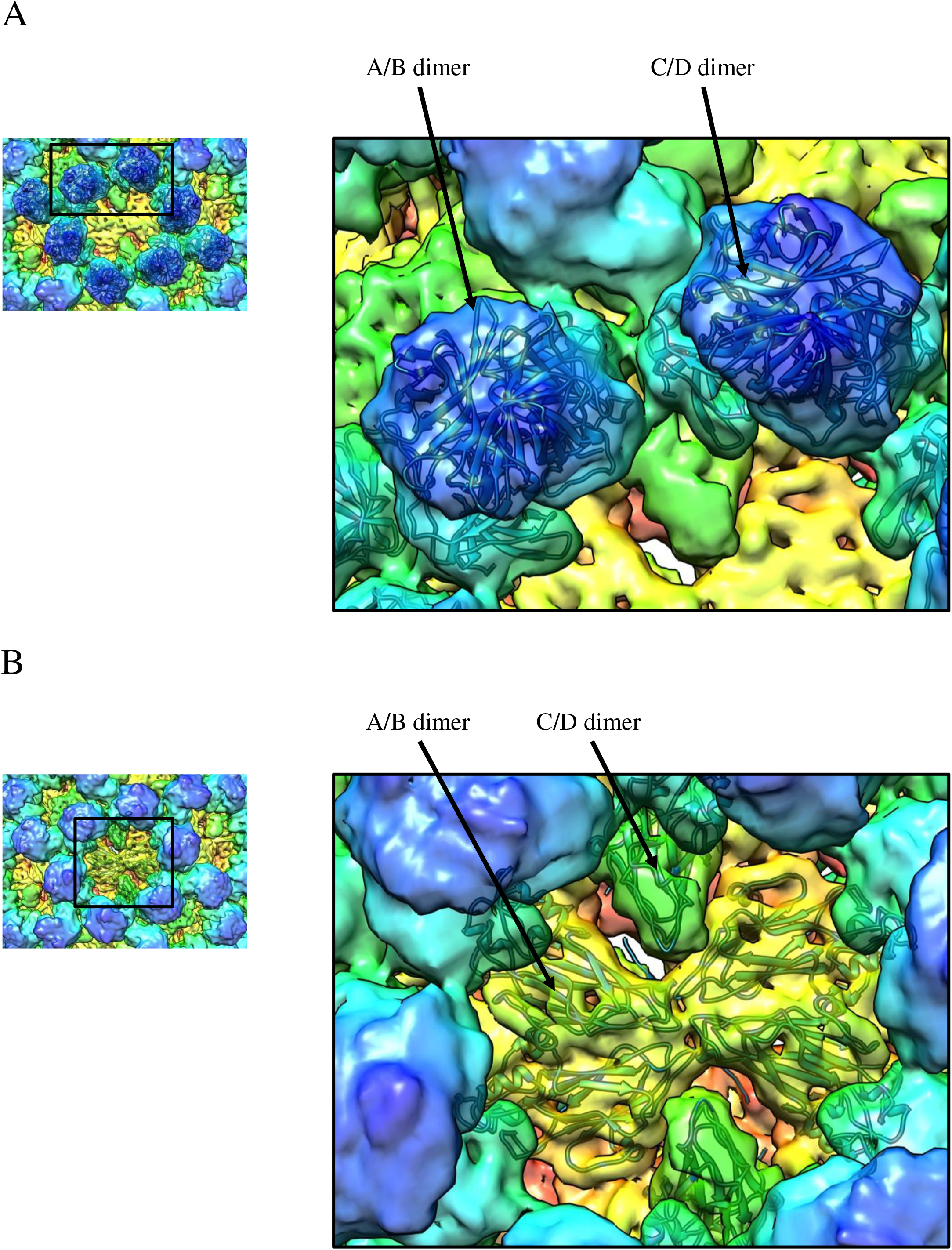
The X-ray crystal structures of NSW-2012 P domain and GI.1 S domain were fitted into the T=4 VLP density map. (A) The X-ray crystal structure of NSW-2012 P domain (4OOS, cartoon) could be fitted into the A/B and C/D P domain dimers, indicating little conformational change. (B) The X-ray crystal structure of the GI.1 S domain (1IHM, cartoon) fitted into the A/B and C/D S domain dimers. However, the cavity and flap-like structures on the D subunit suggests a large conformational change compared to typical T=3 particles.

Unfortunately, it was problematic to fit the hinge region, since the hinge region on the X-ray crystal structure of NSW-2012 P domain was excluded from the expression construct and the hinge region of the GI.1 VLPs was flattened (7, 8). Interestingly, an apparent point of contact was also observed between the S domain and the C-terminus of the P domain on the D subunit (Fig. 8). This may stabilize the convex conformation of the C/D dimer and possibly the T=4 VLPs. Indeed, the C-termini of VP1 on the GI.1 VLPs and the GII.4 P domain were found to be flexible (8, 37). Moreover, the C-terminus of VP1 was previously shown to be important for the size and stability of VLPs (38).

**Figure 8.**
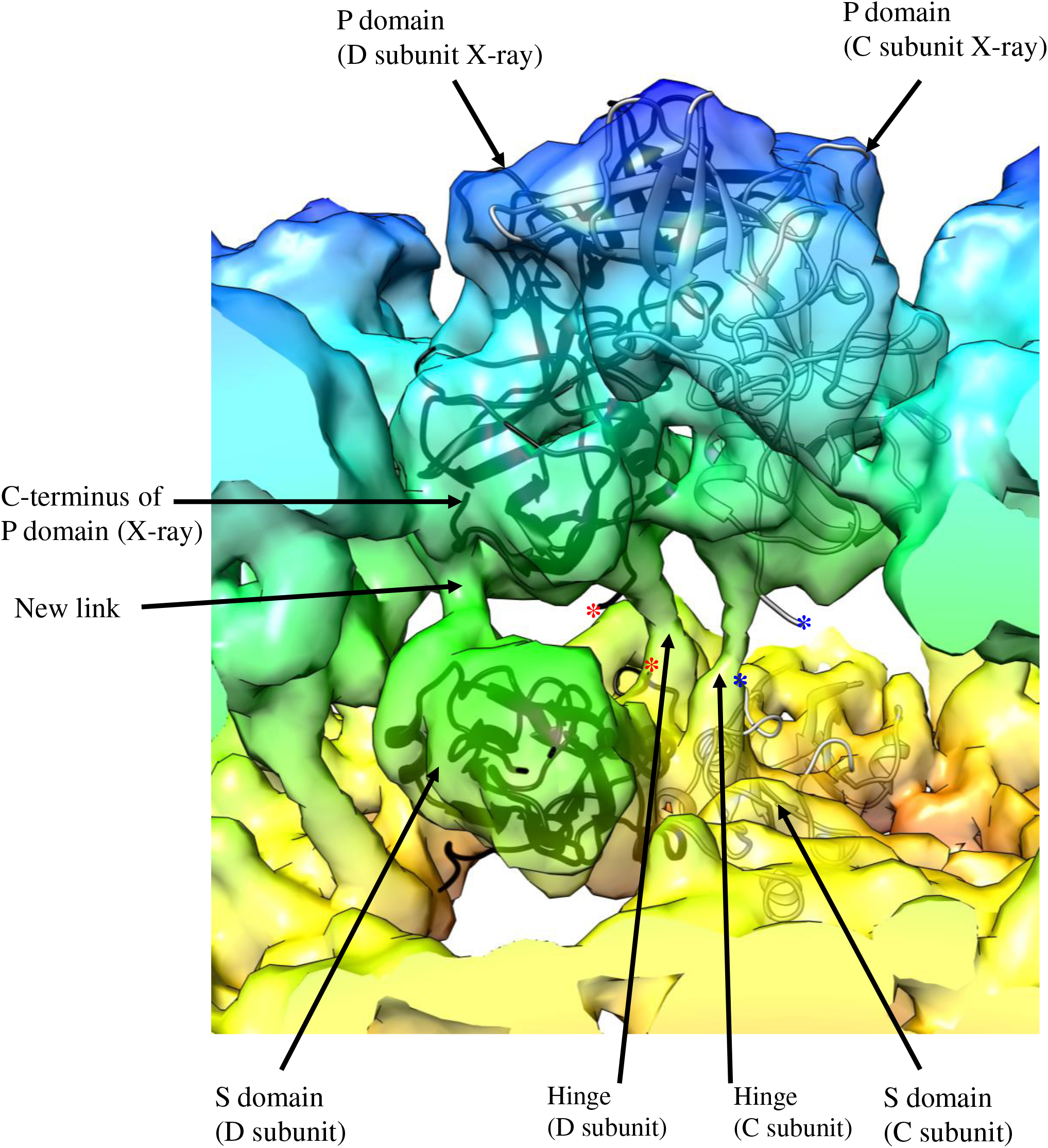
A close-up view of NSW-2012 C/D dimer. The fitted X-ray crystal structures of the GI.1 S domain (cartoon) and the GII.4 P domain (cartoon) into the cryo-EM map shows the how the hinge region connects the S and P domains. Also, the new connection between the S domain and the C-terminus of the P domain is shown. The asterisk represents the missing hinge region on the X-ray crystal structures that connects of the S and P domains for the C subunit (blue) and D subunit (red).

Another interesting feature that we observed with the T=4 VLPs was the raised P domains (Figs. 4 and 8). We found that the T=4 P domain was elevated ∼21-Å off the shell by an extended hinge region, which was higher than the P domains on the GII.10 VLPs, which were raised ∼14-Å (7). The hinge region in NSW-2012 and GII.10 (7) were both ∼10 amino acids and mainly conserved (39). This result suggested that the raised P domains might be a structural feature of GII noroviruses, since the P domains on the GI.1 VLPs were essentially resting on the shell (8).

### Cryo-EM structure of NSW-2012 T=3 VLPs

To determine the structure of a recent GII.4 isolate (NSW-2012), 10,548 particles were imaged by cryo-EM, and a 3D image reconstruction was calculated. Unexpectedly, this analysis revealed a previously unknown T=4 assembly. In this preparation however, a very small number of particles (391 particles) appeared smaller (∼46 nm) than the ∼50 nm VLPs (Fig. 4A). Reconstruction of these particles at ∼15-Å resolution revealed the expected T=3 icosahedral symmetry (Fig. 9). At this resolution, the hinge region was not visible. Also, the cavity and flap-like structures that characterized the T=4 VLP were not present. Based on the T=3 geometry, these VLPs were likely composed of 180 copies of VP1, however the VP1 dimers or subunits were not clearly resolved. Overall, these results indicated that NSW-2012 VP1 formed VLPs of which ∼5% of the population (391 of 10,548) were T=3 particles. This proportion was confirmed over several VLP preparations. Unfortunately, owing to the low numbers it remains challenging to improve on the achieved resolution, since only few T=3 VLPs were expressed and were not able to be separated from the T=4 VLPs using CsCl or sucrose gradient ultracentrifugation.

**Figure 9.**
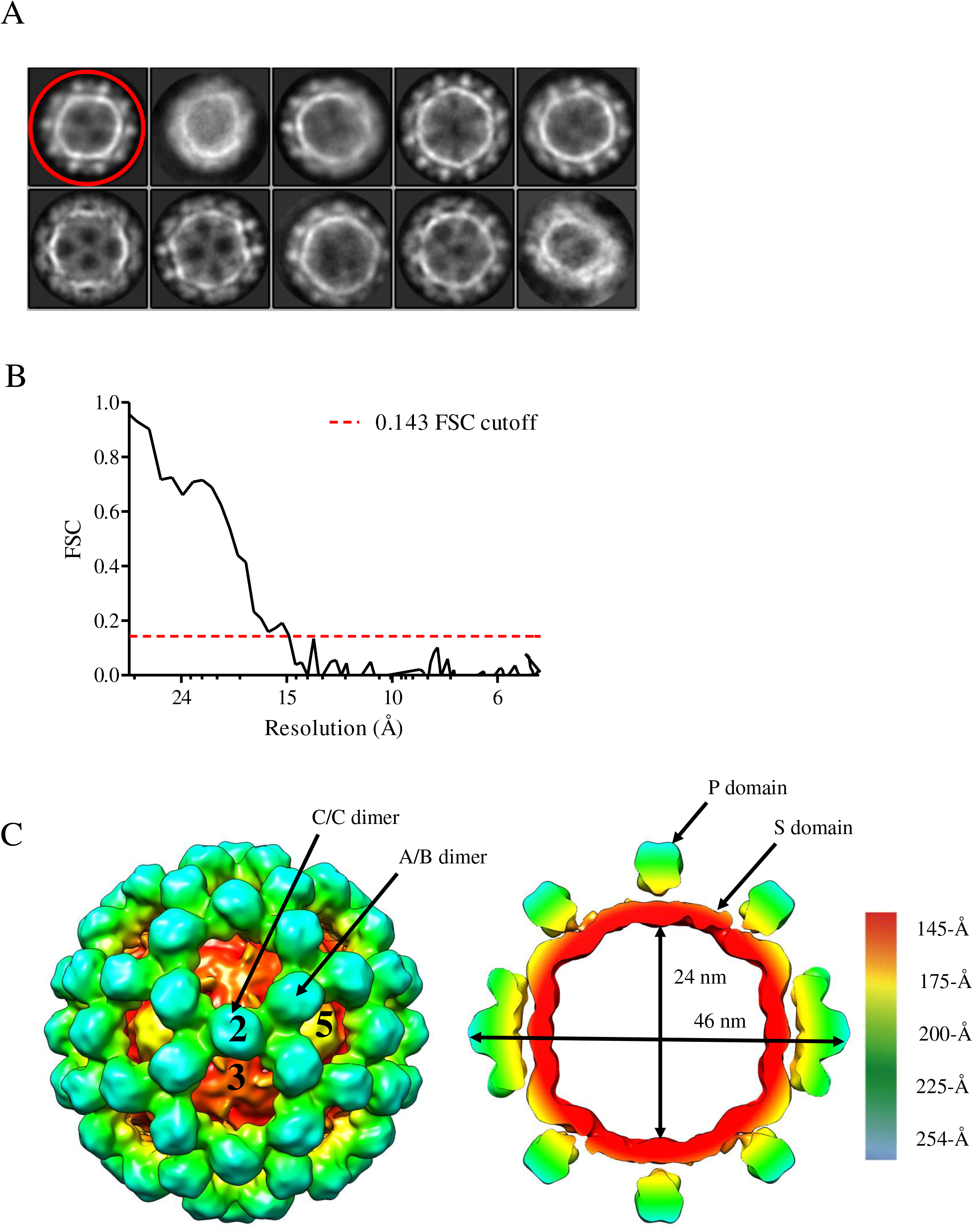
Cryo-EM reconstruction structure of NSW-2012 T=3 VLPs. (A) 2D classification of 1780 manually picked particles. The first class (red circle) shows particles of a smaller diameter, compared to averages of other classes. These smaller particles were used for further refinement. (B) FSC plot of the icosahedral reconstruction of T=3 VLPs indicates a resolution of 15-Å. (C) The left side shows NSW-2012 exhibiting T=3 icosahedral symmetry (symmetry axis labeled 2, 3, and 5). These VLPs were composed of 180 copies of VP1 that forms A/B and C/C dimers. The cutaway section (right) shows the inner and outer diameter of the particle, which measured 24 and 46 nm, respectively. At this resolution the hinge region could not be resolved, but the large gap between S and P domain indicates that the P domains are raised up from the shell.

### Cryo-EM structure of GII.4 CHDC-1974 T=4 VLPs

To test whether assembly of VLPs with T=4 icosahedral symmetry is a property of VP1 from all GII.4 strains, we proceeded to determine the cryo-EM structure of VLPs produced by VP1 of CHDC-1974. The VLPs were mostly mono-disperse and homogenous in size (Fig. 10A). From 591 images, 42,485 particles were processed to calculate a reconstruction with a final resolution of 6.1-Å (Fig. 10B). The CHDC-1974 VLPs also had T=4 symmetry (Fig. 11) and their structure closely resembled that of NSW-2012 VLPs. Unlike NSW-2012 however, CHDC-1974 VP1 only assembled into T=4 VLPs and no T=3 VLPs were identified.

**Figure 10.**
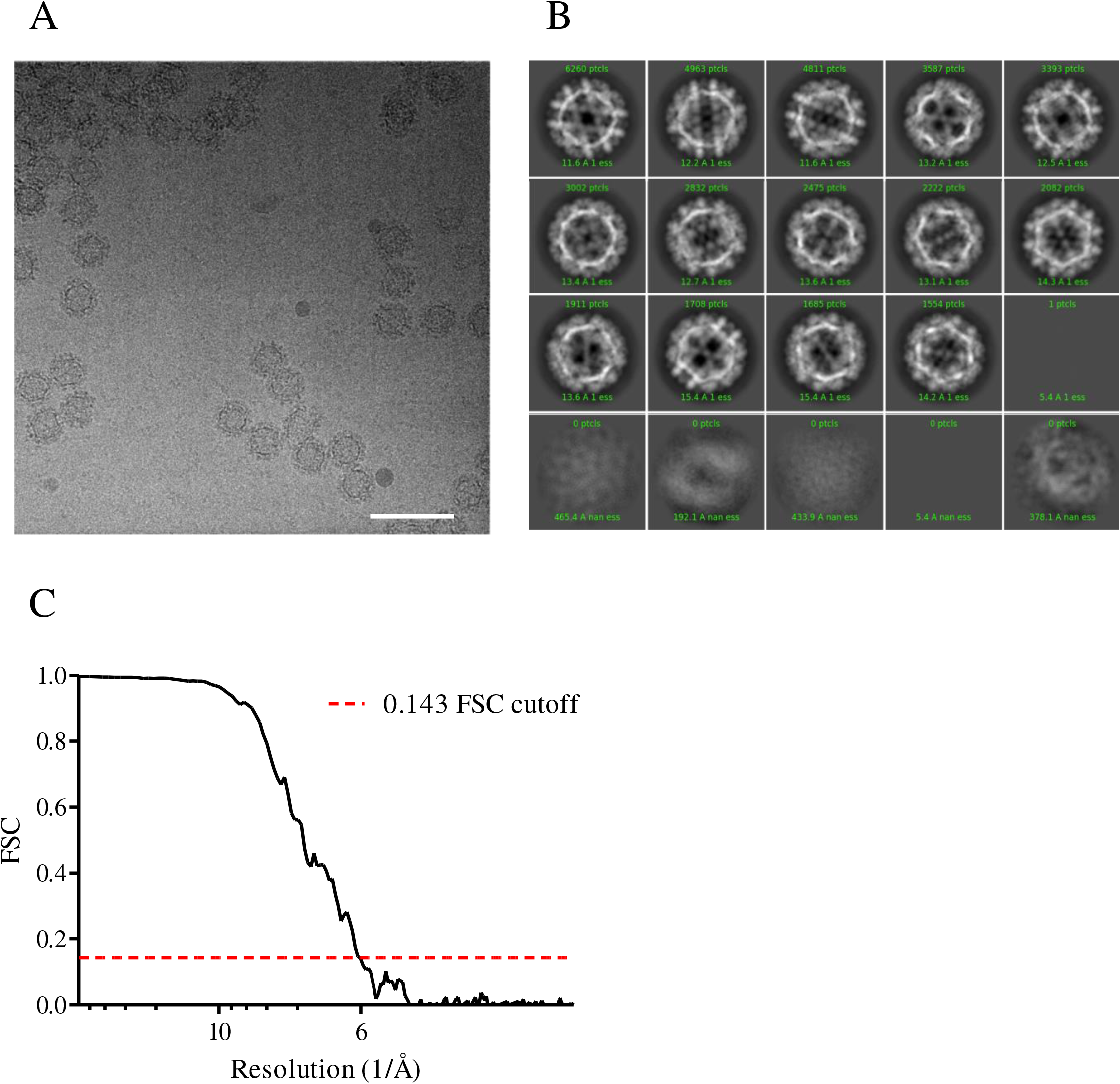
CHDC-1974 cryo-EM data processing. (A) A representative cryo-EM micrograph of CHDC-1974 VLPs at 64,000× magnification. The scale bar represents 100 nm. (B) 2D classification of CHDC-1974 particles. (C) FSC plot of the icosahedral reconstruction of CHDC-1974 indicates a resolution of 6.1-Å at 0.143 cutoff.

**Figure 11.**
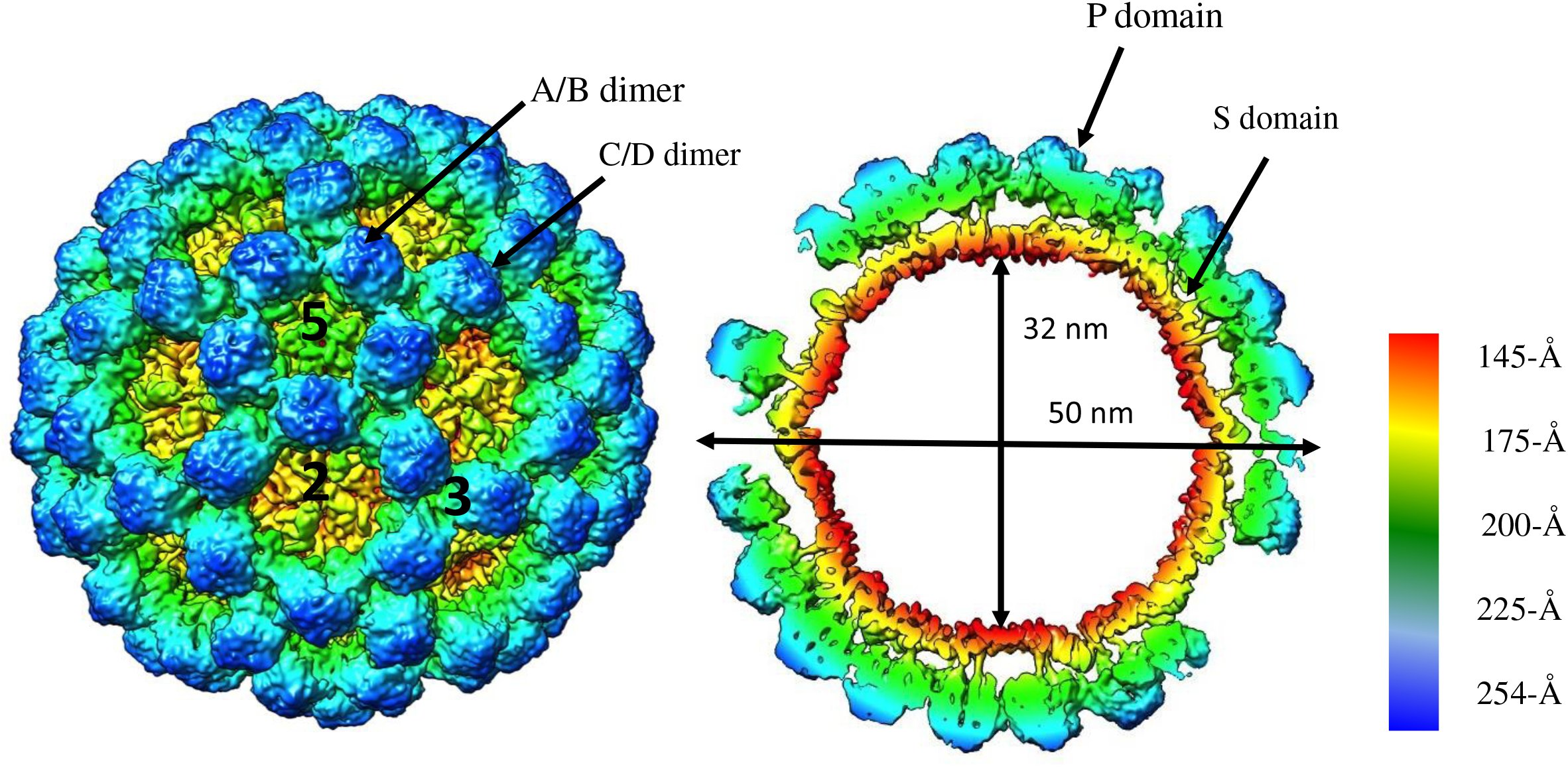
Cryo-EM structure and analysis of CHDC-1974 VLPs. The image on the left side shows that CHDC-1974 VLPs has a T=4 icosahedral symmetry (symmetry axis labeled 2, 3, and 5) and was composed of 240 copies of VP1. The VP1 exhibited four quasiequivalent conformations (A, B, C, and D) that gave rise to two distinct dimers (A/B and C/D). At the icosahedral twofold axis, the B, C, and D subunits were alternating, while the A subunit was located around the fivefold axis. The right side shows a cutaway section of these VLPs and indicates that the inner and outer diameters are 32 nm and 50 nm, respectively. The P domains are elevated ∼21 Å off the S domain.

We found that CHDC-1974 T=4 VLPs were also composed of 240 copies of VP1 that formed the quasi-equivalent subunits A, B, C, and D and A/B and C/D dimeric capsomeres. The inner diameter of the shell was 32 nm, whereas the outer diameter of the capsid was 50 nm. As in NSW-2012 VLPs, cavity and flap-like structures were also present on the CHDC-1974 VLPs (Fig. 12). The CHDC-1974 A/B and C/D dimers showed a similar convex conformation as NSW-2012 dimers, although slightly less pronounced (Fig. 13). The X-ray crystal structure of CHDC-1974 P domain (5IYN) was easily fitted into the CHDC-1974 VLP density map (Fig. 14A). The GI.1 S domain also fitted into the A, B, and C subunits, whereas the GI.1 S domain was repositioned to occupy the D subunit density (Fig. 14B). Similar to NSW-2012 VLPs, a possible contact interface was observed between the S and P domains on the D subunit (Fig. 15). Lastly, we found that CHDC-1974 P domain was also lifted off the shell ∼21-Å by an extended hinge region (Figs. 11 and 15).

**Figure 12.**
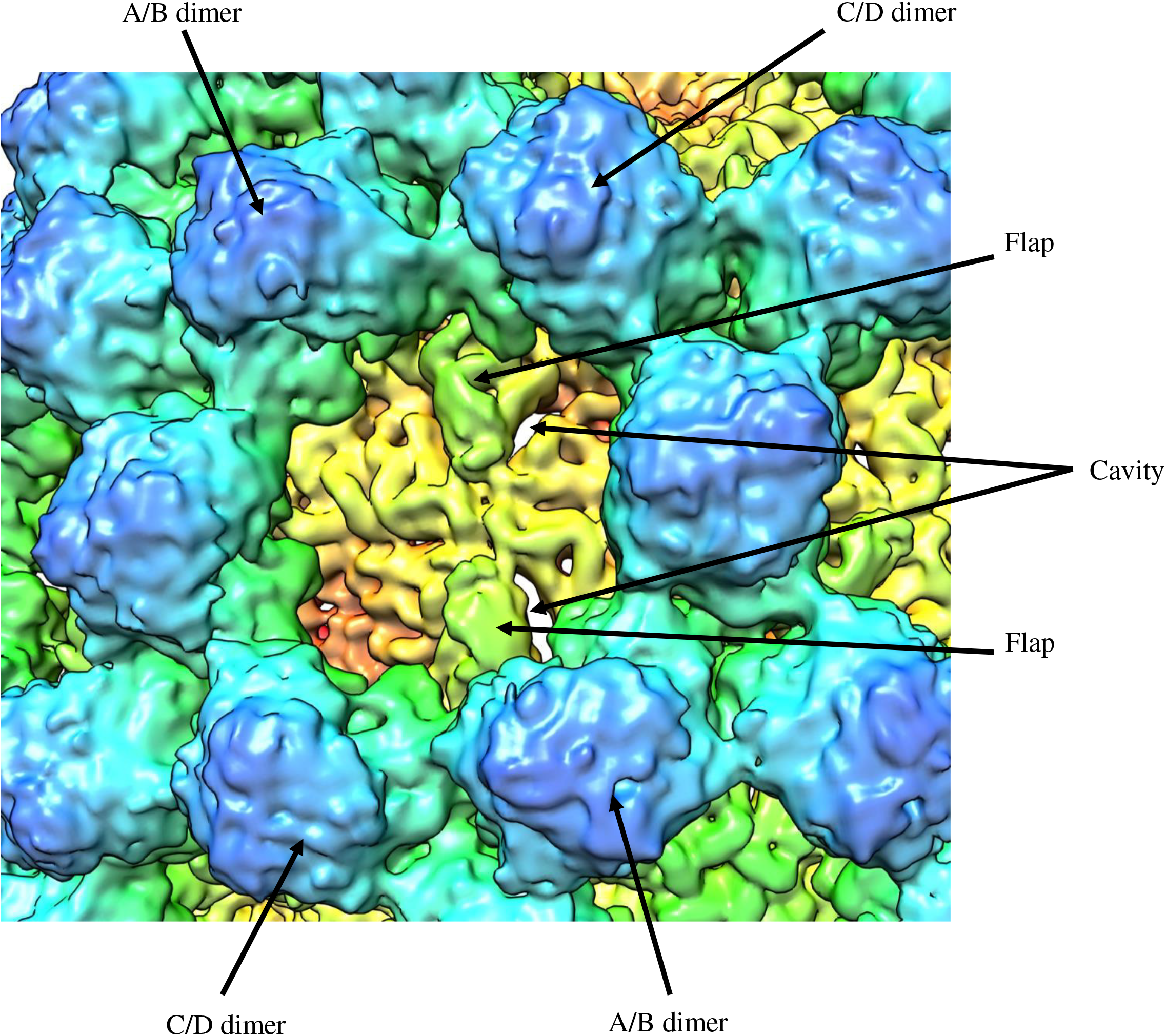
CHDC-1974 T=4 VLPs shows several new structural features. The cavity and flap-like structures are observed at the twofold axis and are found on opposing sides. The cavity and flap-like structures are associated with the S domain on the D subunit.

**Figure 13.**
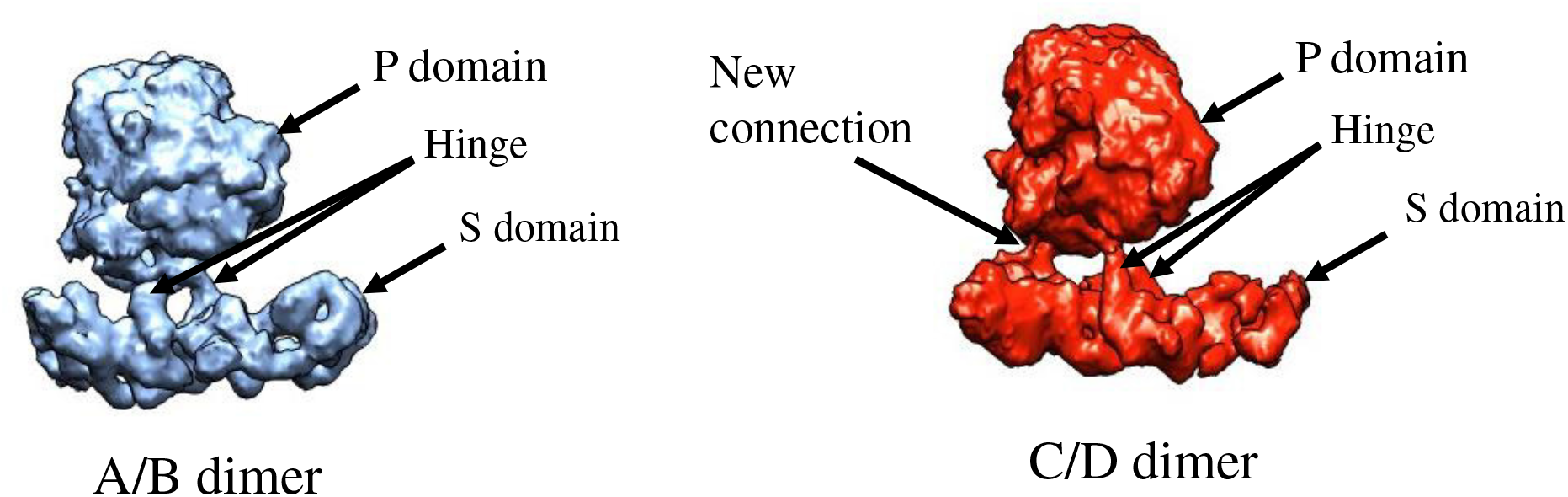
CHDC-1974 T=4 VLPs are formed with 60 A/B and 60 C/D VP1 dimers. The A/B and C/D dimers show an equivalent convex confirmation on the S domain. Also, the additional connection between the D subunit of the S and P domain was found on the CHDC-1974 T=4 VLPs.

**Figure 14.**
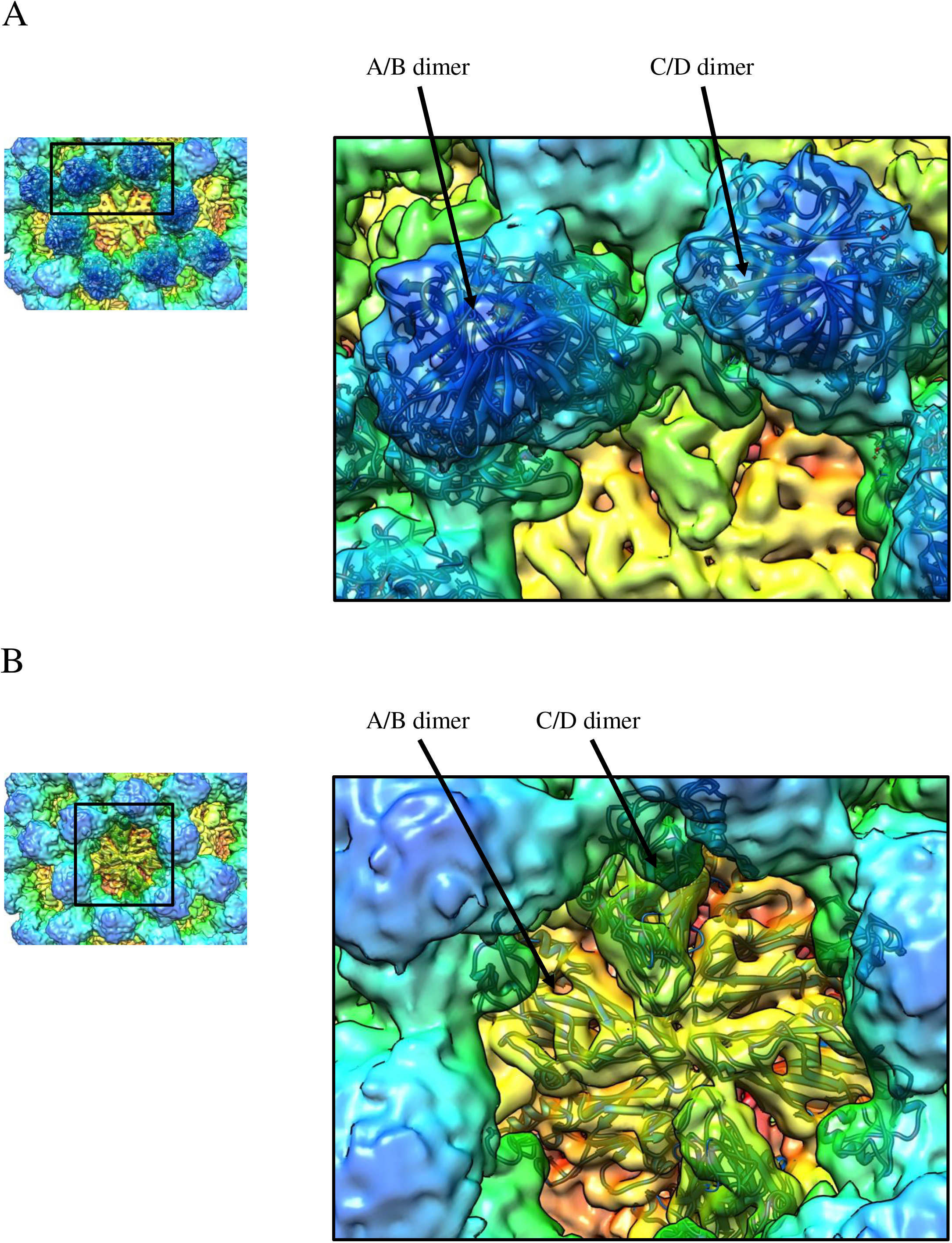
The X-ray crystal structures of CHDC-1974 P domain and GI.1 S domain were fitted into the VLP density map. (A) The X-ray crystal structure of NSW-2012 P domain (5IYN, cartoon) easily fitted into the A/B and C/D P domain dimer densities. (B) The X-ray crystal structure of the GI.1 S domain (1IHM, cartoon) fitted into the A/B and C/D S domain dimers. However, the cavity and flap-like structures on the D subunit suggests a large conformational change from typical T=3 particles.

**Figure 15.**
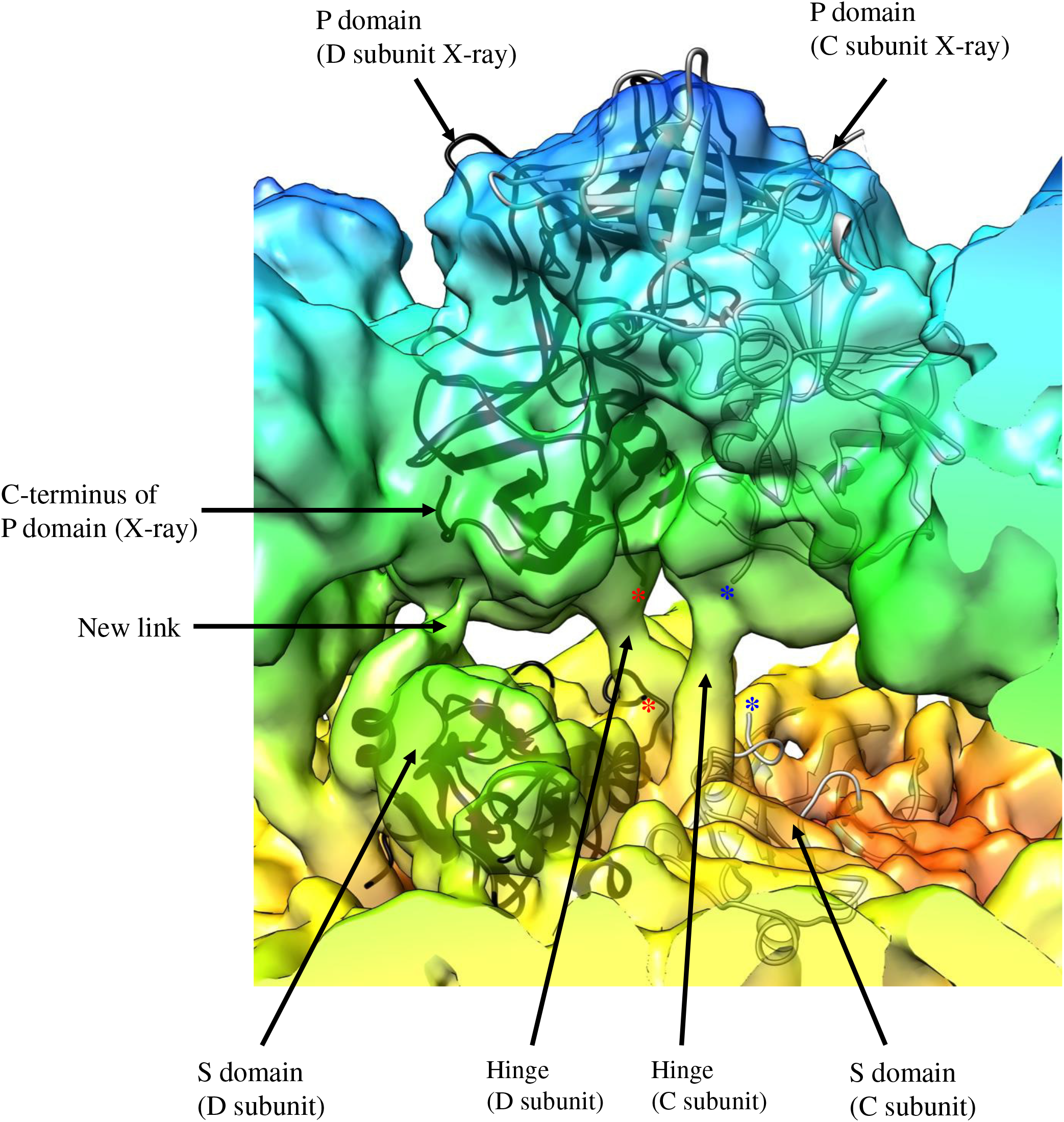
A close-up view of CHDC-1974 C/D dimer. The fitted X-ray crystal structures of the GI.1 S domain (cartoon) and the GII.4 P domain (cartoon) into the cryo-EM map shows the how the hinge region connects the S and P domains. Also, the new connection between the S domain and the C-terminus of the P domain is shown. The asterisk represents the missing hinge region on the X-ray crystal structures that connects of the S and P domains for the C subunit (blue) and D subunit (red).

Overall, these results showed that GII.4 VP1 sequences isolated over three decades apart remained structurally conserved. This could imply that other GII.4 VP1 sequences would also form T=4 VLPs when expressed in insect cells, especially since these two sequences had only 89% amino acid identity.

### Previous binding studies using GII.4 capsids

GII.4 VLPs and their corresponding P domains have been extensively examined for binding to co-factors (HBGAs and bile acid), antibodies, human milk oligosaccharides (HMOs), and Nanobodies (21, 31, 37, 39-41). NSW-2012 P domains were capable of binding numerous HBGA types and the VLPs cross-reacted with Nanobodies and antibodies that were raised against other genotypes or GII.4 variants. However, the NSW-2012 VLPs did not bind bile acid and poorly bound HMOs (unpublished). Overall, these results suggested that T=4 VLPs were indeed antigenically relevant and capable of functioning similarly to GII.4 virions with respect to antigenicity and HBGA interactions.

### GII.4 VLPs in vaccine trials

VLP vaccines against norovirus that are currently in clinical trials use a combination of GI.1 VLPs and modified GII.4 VLPs (termed GII.4c) (42, 43) Negative stain EM images of these GII.4c VLPs showed the typical norovirus morphology. However, the size determination and the structure are not available. Interestingly, GII.4c and NSW-2012 shared 94% amino acid identity with most substitutions (28 of 31) located in the P domain (Fig. 1). Therefore, it is tempting to speculate that GII.4c VP1, which closely matched NSW-2012 VP1 sequence, might also form T=4 VLPs particles. In both VLPs, VP2 is not present.

To test whether GII.4 virions are T=3 or T=4 assemblies, we used negative stain EM of authentic GII.4 virions. These virion images revealed a smaller diameter of ∼44 nm compared to the T=4 VLPs (∼50 nm) (Fig. 16). This size corresponds to the diameter determined for the NSW-2012 T=3 VLPs and indicates that GII.4 virions likely exhibit T=3 icosahedral symmetry.

**Figure 16.**
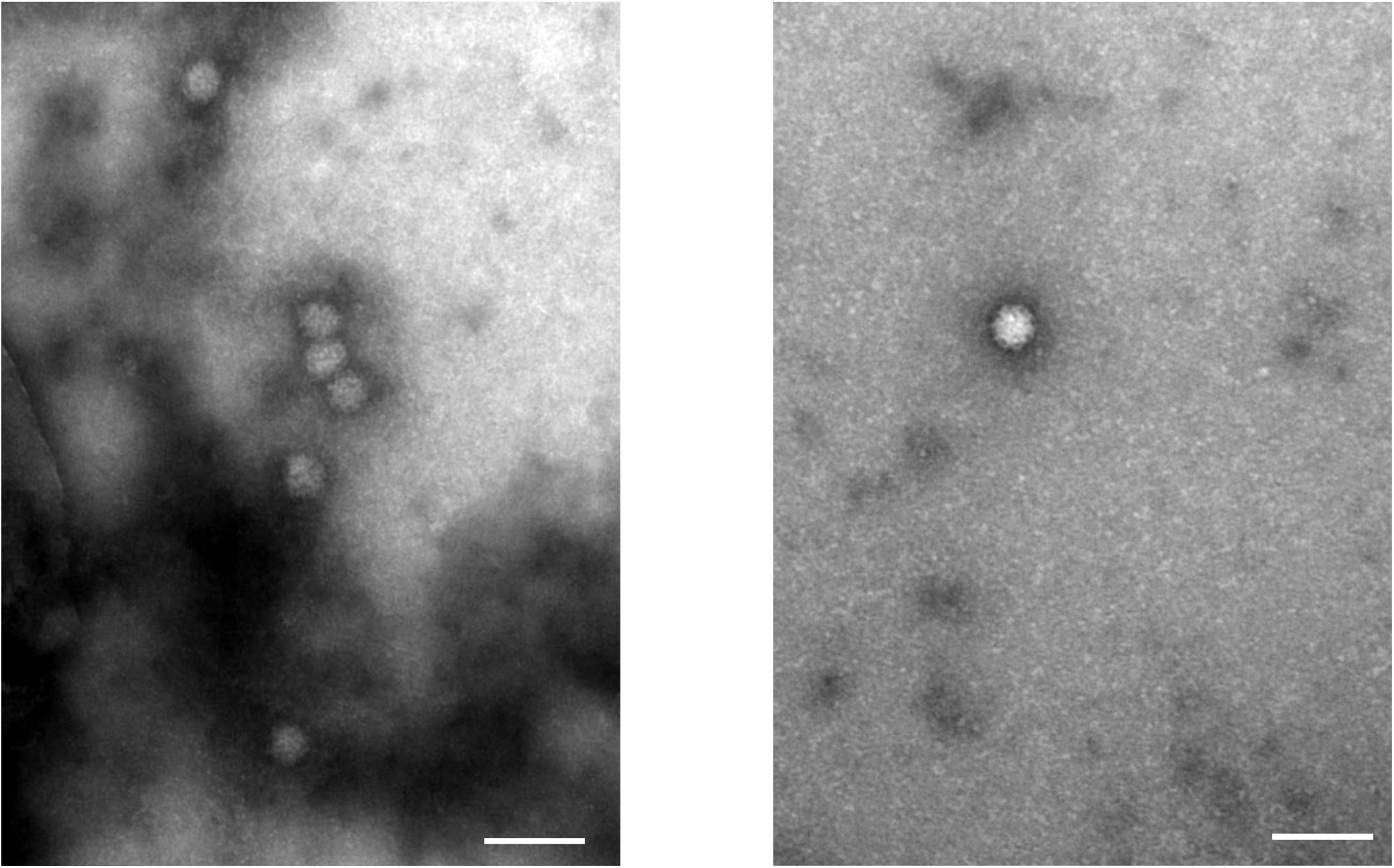
EM images of GII.4 virions. Negative stain EM images of GII.4 virions show that the virions exhibit a smaller diameter (∼44 nm) than GII.4 VLPs expressed in insect cells that were ∼52 nm (see Figure 2).

In general, studies have shown that norovirus-specific antibody titers were raised after vaccination with VLPs, but the levels of protection were not strongly improved compared to placebo groups (44). This might indicate that vaccination leads to production of neutralizing antibodies against epitopes on T=4 VLPs that are not accessible on T=3 virions (i.e., when challenged) and that therefore vaccine efficacy could be lowered by a sub-optimal antigen. Clearly, further studies are needed in order to determine whether the GII.4c VLPs form T=4 particles and if the T=4 VLPs are antigenically identical to T=3 VLPs.

### Summary

We have shown that upon heterologous expression of human norovirus GII.4 VP1 in insect cells, VLPs are formed that adopt T=4 icosahedral symmetry. This is at odds with the likely T=3 symmetry virions encoded by this virus. There are two important consequences of this outcome. Firstly, the assembly of GII.4 T=4 VLPs may impact on results from previous studies in which it was assumed that GII.4 norovirus VLPs were morphologically similar to virions. Secondly, the cavity and flap-like structures on the T=4 could elicit antibodies that are not capable of recognizing T=3 virions. Further structural and binding studies using a library of GII.4-specific Nanobodies are planned in order to investigate these novel epitopes.

## ACKNOWLEDGEMENTS

We acknowledge the excellence cluster CellNetworks (Cryo-EM network) of the University of Heidelberg for cryo-EM data collection, the EM core facility at DKFZ, and Baden-Württemberg High Performance Cluster (bwHPC). We thank Anna Koromyslova for EM images of GII.4 virions and Benedikt Wimmer for setting up the cryo-EM software. The funding for this study was provided by the CHS foundation; the Baden-Württemberg Stiftung (GLYCAN-BASED ANTIVIRAL AGENTS); Deutsche Forschungsgemeinschaft (DFG, FOR2327); and the BMBF VIP+ (Federal Ministry of Education and Research) (NATION, 03VP00912).

